# Detection and characterization of circulating microvesicles containing Shiga toxin type 2 (Stx2) in a rat model of Hemolytic Uremic Syndrome

**DOI:** 10.1101/2022.02.17.480856

**Authors:** Flavia Sacerdoti, Fernando Gomez, Carolina Jancic, Marcela A. Moretton, Diego A. Chiappetta, Cristina Ibarra, María Marta Amaral

## Abstract

Shiga toxin (Stx) producing *Escherichia coli* (STEC) are foodborne pathogens that release Stx and may develop Hemolytic Uremic Syndrome (HUS). Stx causes endothelial cell damage and leads to platelets deposition and thrombi formation within the microvasculature. It has been described that Stx activates blood cells and induces the shedding of proinflammatory and prothrombotic microvesicles (MVs) containing the toxin. In this sense, it has been postulated that MVs containing Stx2 (MVs-Stx2+) can contribute to the physiopathology of HUS, allowing Stx to reach the target organs and evading the immune system. In this work, we propose that circulating MVs-Stx2+ can be a potential biomarker for the diagnosis and prognosis of STEC infections and HUS progression. In this regard, we developed a rat HUS model by the intreperitoneal injection of a sublethal dose of Stx2 and observed: decrease in body weight, increase of creatinine and urea levels, decrease of creatinine clearance and histological renal damages. After characterization of renal damages we investigated circulating total MVs and MVs-Stx2+ by flow cytometry at different times after Stx2 injection. Additionally, we evaluated the correlation of biochemical parameters such as creatinine and urea in plasma with MVs-Stx2+. As a result, we found a significant circulation of Mvs-Stx2+ at 96 hours after Stx2 injection, nevertheless no correlation with creatinine and urea plasma levels were detected. Our results suggest that MVs-Stx2+ may be an additional biomarker for the characterization and diagnosis of HUS progression. Further analysis is required in order to validate MVs-Stx2+ as biomarker of the disease.

## INTRODUCTION

The main cause of Hemolytic Uremic Syndrome (HUS) in children is Shiga toxin (Stx)-producing *Escherichia coli* (STEC) infection. HUS is a disease clinically characterized by microangiopathic hemolytic anemia, thrombocytopenia, and acute kidney injury [1]. Argentina has the highest incidence rate of HUS in the world due to STEC infections. The most accepted explanation about this epidemiological situation is the concomitance of different factors as: contamination of food or water, person-to-person transmission, poor hygiene practices, and the circulation of more virulent strains [2]. According with the National Health Surveillance System, in 2019 343 new cases of HUS and 305 in 2020 were notified [3]. Also, in this country, typical HUS or associated with STEC infections is the principal cause of acute renal injury in pediatric age groups and the second most recurrent cause of chronic renal disease [4, 5]. STEC can colonize the intestine where Stx is released and after intestinal damage, the toxin can access the bloodstream [6]. In circulation, Stx induces endothelial cell damage and as consequence, platelet deposition and thrombi formation within the microvasculature is produced, inducing thrombotic microangiopathy that affect mostly the kidney. In this regard, Stx may circulate in the free form or bound to blood cells such as red blood cells, neutrophils and platelets [7]. In this sense, it has been reported that Stx activates blood cells that induce the shedding of proinflammatory and prothrombotic microvesicles (MVs) containing Stx that may in this way reach the target organs.

MVs are small plasma membrane structures sizing from ∼100-1000 nm, shared by cells upon activation, stress and apoptosis [8-10]. MVs can act as important signaling structures and mediate the communication between vascular cells allowing membrane interactions between distant cells. MVs released may also maintain cellular integrity in order to remove harmful substances and eliminate unwanted components. Circulating levels of MVs are altered during several diseases such as autoimmune diseases [11] coagulation disorders [12], cardiovascular diseases [13], cancer [14] and infections [15], among others. MVs can be detected in peripheral blood in healthy conditions and may be originated by different blood cell types including platelets, endothelial cells, erythrocytes and granulocytes. Moreover, membrane of MVs may be enriched in proteins and receptors derived from their parent cells, and intravesicularly may transport mRNA, enzymes and transcription factors and others [16]. MVs may show procoagulant properties probably induced by phosphatidylserine (PS) exposure and the presence of tissue factor on their membrane. So, high circulating levels of MVs can be deleterious and may trigger procoagulant and inflammatory profiles that may lead to adverse clinical outcome [17]. On the other hand, MVs are proposed as potential biomarkers for the diagnosis and prognosis of vascular diseases and complications, and the assessment of treatment response [18].

Futhermore it has been proposed that MVs shed by blood cells during STEC infection may participate in the prothrombotic lesion, hemolysis and the transfer of the toxin from circulation into the cells of the target organs, mainly the kidney [19, 20]. Thus, MVs transporting Stx have been suggested as an important mechanism for Stx inducing systemic and target injury [21]. Interestingly, the transport of Stx in circulating MVs enables toxin evasion of the immune system and protection from degradation, by the way they may contribute to the disease and may be considered as potential biomarkers. Considering these backgrounds, the aim of this work was to analyze the presence of blood circulating MVs-Stx2+ in a rat model of sublethal HUS, in attempt to introduce it detection as a new clinical biomaker for HUS early diagnosis.

## MATERIALS AND METHODS

### Drugs and Chemicals

Purified Shiga toxin type 2a (Stx2) was purchased from Phoenix Laboratory (Tutfs Medical Center, Boston, MA, USA) and was checked for LPS contamination by the *Limulus amoebocyte lysate assay* (LAL test), Biowhittaker Inc. (Marylans, USA). Toxin was diluted with sterile phosphate buffered saline (PBS) before injection and the final solution contained < 10 pg LPS/ng of pure Stx2.

### Animals

Sprague Dawley adult female rats (180-250 g; three months of life) were acquired from the Animal facility of the School of Pharmacy and Biochemistry-University of Buenos Aires. Animals received food and water *ad libitum* and were housed under controlled conditions of light (12-h light, 12-h dark) and temperature (23-25°C). This study was carried out in strict accordance with the recommendations detailed in the guide for the care and use of Laboratory Animals of the National Institutes of Health. Protocols were approved by the Committee for the Care and Use of Laboratory Animals of the School of Medicine of the University of Buenos Aires (CICUAL, Permit Number 2439/2019).

### Animal Protocol

Two groups of rats were injected with a sublethal dose of Stx2 (0.5 ng/μl/g of body weight total, n=6) or PBS (1 μl/g of body weight total, n=4). After injection, the animals were housed as mentioned before and weighed daily during the first four days and at day nine. Data of weight are presented as Δ weight - (body weight at n day) (body weight at 0 days) after injection. Additionally, a blood sample from the tail vein was collected in a tube with sodium citrate (3.8 %) for analysis of plasma urea, plasma creatinine and microvesicles (MVs) isolation, every 24 hs for four days. The experiment was repeated three times. On the other hand, four days after treatment, another group of rats (Stx2 n=6 and PBS n=4) were anesthetized intraperitoneally with 100 μg ketamine and 10 μg diazepam per g of body weight. Animals were perfused firstly with a 0.9% NaCl solution (w/v) and then with 4% (v/v) formaldehyde in PBS. Finally, the kidneys were removed for immunohistochemical evaluation. In addition, another group of four Control and four Stx2 injected rats were placed in metabolic cages for 24 h (between 48 h and 72 h after injection) for food intake, urine volume determination and creatinine clearance (CrCl) determination. Additionally, body weight and ingestion of food were analyzed. The experiments were repeated twice (**Figure 1, Flow chart protocol**).

**Figure 1.**
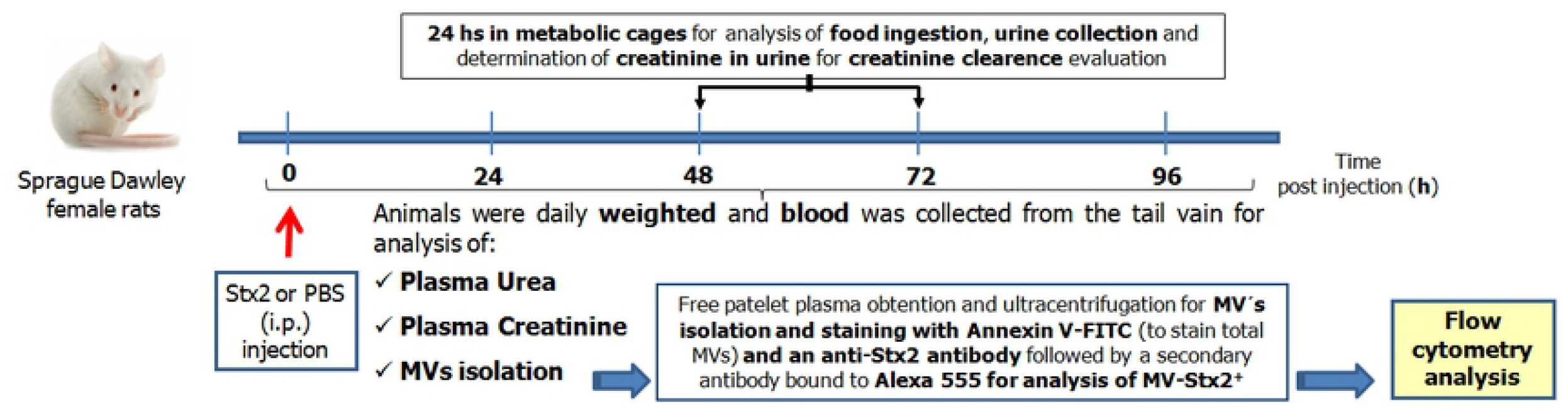
Flow chart of protocol used in Stx2 and PBS injected rats. Rats were i.p injected (t=0) with a sublethal dose of Stx2 (n=6) or PBS (n=4) weighted daily and blood was collected at different times (0, 24, 48, 72 and 96 h after injection) for the evaluation of: plasma urea, plasma creatinine and MVs isolation. Another group of rats were housed in metabolic cages from 48 to 72 h after injection for evaluation of: food ingestion, urine collection and creatinine determination for the evaluation of renal function by creatinine clearance. Total MVs were isolated from plasma and MVs containing Stx2 (MV-Stx2+) were analyzed by Flow Cytometry.

### Plasma urea and creatinine evaluation

Plasma collected at 48, 72 and 96 h after Stx2 or PBS injections were evaluated for urea and creatinine levels. Urea and creatinine concentration (mg/dL) were determined by using commercial kits (Kinetic Creatinine AA and Urea color 2R, Wiener Lab, Argentina).

### Histological evaluation of renal damages

Rat kidneys were fixed for 48 h with formol 10% in PBS 0.1M (pH 7.4), dehydrated, and embedded in paraffin. Sections of 5 μm were made using a microtome (Leica RM 2125, Wetzlar, Germany) and mounted on 2% silane coated slides. The sections were stained with Periodic Acid-Schiff (PAS) and then observed and photographed under light microscopy (Nikon Eclipse 200, Melville, NY, USA). Three randomly selected images from the cortex of each slide were obtained. Renal damage was evaluated by semi-quantitative scoring of tubular necrosis (% of necrotic tubes/total tubes) that was characterized by loss of the brush border, flattening of cells, rupture, and detachment of tubular cells from the basement membranes. The total percentage of damage for each treatment was calculated from the average of the scored images from each group. The results were expressed as the mean ± SEM of the percentage of tubular necrosis.

### Obtention of microvesicles (MVs)

MVs were obtained according to the protocol previously described by Stahl *et al* [20], with minor modifications. Firstly, blood samples were centrifuged for 15 min at 600 *g* at 20°C to remove cells and to obtain plasma fraction. Then, plasma was centrifuged at 13000 *g* for 3 min to obtain platelet-free-plasma (PFP). Then, PFP was centrifuged at 23000 *g* at 15°C for 45 min to obtain the fraction of MVs.

### Evaluation of the hydrodynamic diameter and size distribution of MVs by Dynamic Light Scattering (DLS)

MVs size distribution was measured by dynamic light scattering (DLS, scattering angle of θ= 173° to the incident beam, Zetasizer Nano-ZSP, ZEN5600, Malvern Instruments, United Kingdom) at 37°C. Briefly, fifty microliters of isolated MVs were dispersed in 1 ml of Phosphate buffer. Then aliquots (1 ml) were vortexed for 30 seconds and samples were equilibrated over 5 minutes before each measurement. Results were expressed as mean ± standard deviation (SD, n = 5).

### Total MVs and Stx2 containing MVs (MVs-Stx2+) detection and analysis

Obtained MVs, from blood at 24, 48, 72 and 96 h after Stx2 injection or Control, were fixed with paraformaldehyde 1% (PFA, Sigma Aldrich, USA). Then, MVs were labeled with Annexin V-FITC (AV-FITC, Sigma Aldrich, USA) and quantified by Flow cytometry at 48, 72 and 96 h for Stx2 and Control rats. Additionally, MVs containing Stx2 were evaluated. For that purpose, MVs were permeabilized with saponin 0.1% (Sigma Aldrich, USA) in Binding Buffer (BB: HEPES-NaOH 0.1 M; NaCl 1.4 M; CaCl_2_ 0.025 M; pH: 7.5). After that, they were incubated with a primary anti-Stx2 antibody (MabC11, kindly provided by Dr. Roxane Piazza [22] for 90 min at room temperature. After incubation MVs were diluted with BB and centrifuged at 21000 *g* for 45 min. Afterward, MVs were resuspended in 100 μl and incubated with a secondary antibody (anti mouse IgG - Alexa 555, Becton Dickenson) for 1 h at room temperature. Finally, MVs were stained with AV-FITC and samples were analyzed by Flow cytometry (PartecX-PASIIIX). MVs were identified as single positive population for AV-FITC and MVs-Stx2+ population as double positive for Annexin V-FITC and Alexa 555. In order to detect Stx2 on the surface of MVs, some experiments were carried out without saponin.

### Data Analysis

Data in images are presented as mean ± SEM. Statistical analysis was performed using GraphPad Prism Software 5.0 (San Diego, CA, USA). ANOVA was used to calculate differences between groups and Tukey’s multiple comparisons test was used as a posttest. When two groups were compared, a non parametric t-test was used. Correlation and linear regression analysis were performed to evaluate the possible association between urea or creatinine and MVs-Stx2+ levels 96h after injection.

## RESULTS

### Body weight and food ingestion after Stx2 injection

Rats treated with Stx2 showed a significant decrease in total body weight at 3, 4 and 9 days post injection compared to Controls (**Figure 2A**). Additionally showed a general discomfort, quiescence and in some cases piloerection. A peak in body weight loss was observed at four days after toxin exposure, then rats recovered weight gradually but did not reach it as controls. On the contrary, Control rats had a gradual increase in total body. Concomitantly, food ingestion, evaluated at 48 h after toxin injection, was significantly lower in Stx2 treated rats compared to Controls (**Figure 2B**).

**Figure 2.**
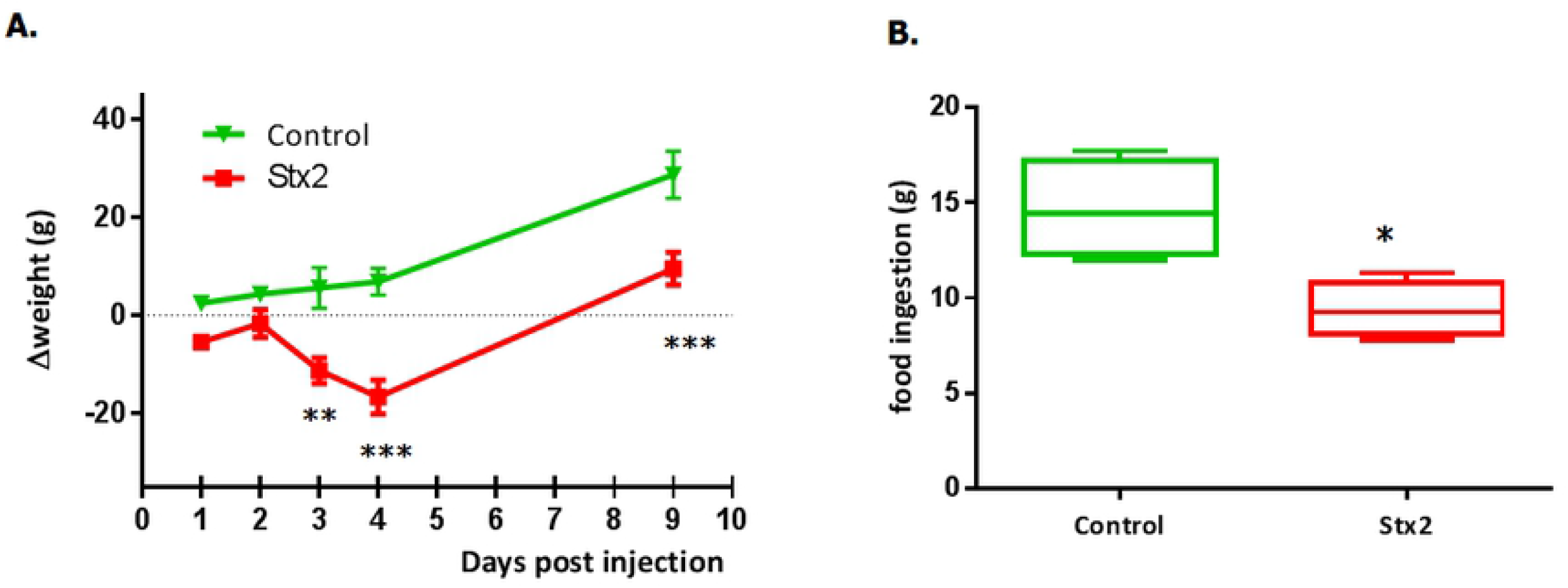
Body weight progression and food ingestion after Stx2 injection. Rats were i.p injected with a sublethal dose of Stx2 (n=6) or PBS (Control, n=4) and body weight was registered (A). Animals were housed, 48 h after Sx2 (n=4) or PBS injection (Control, n=4), in metabolic cages for 24 h and food ingestion was evaluated (B). Data are represented as mean ± SEM. *P<0.05, **P<0.01 and ***P<0.001.

### Evaluation of creatinine and urea in plasma and creatinine clearance

Creatinine (**Figure 3A**) and urea (**Figure 3B**) were evaluated in plasma at 48, 72 and 96 h after Stx2 injection. A statistically significant increase in plasma creatinine was observed at 72 and 96 h after toxin exposure compared to Controls. Additionally, an increase in plasma urea levels was observed at 96 h after Stx2 injection. Also, after 72 h of treatment, a significant decrease in the creatinine clearance was observed in Stx2 injected rats (**Figure 3C**).

**Figure 3.**
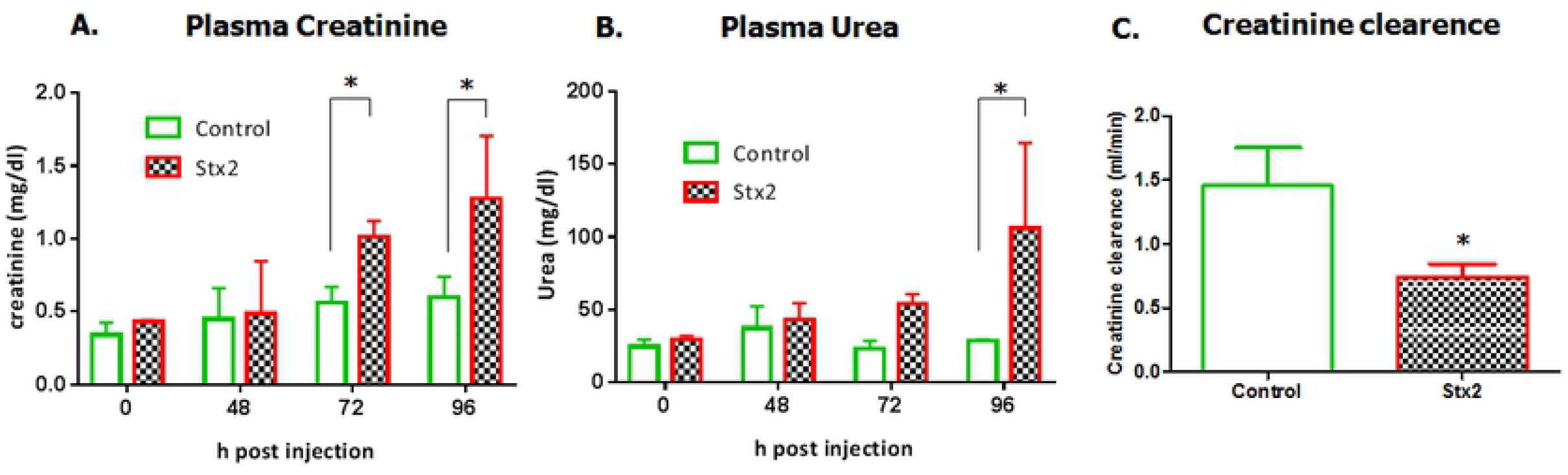
Plasma creatinine, Plasma urea, and creatinine clearence after Stx2 injection. Plasma creatinine (A) and urea (B) were evaluated at different times (0, 48, 72 and 96 h) after Stx2 (n=4) or PBS (Control, n=4) injection. After 48 h of toxin injection, urine from animals housed in metabolic cages was collected for 24 h and creatinine clearence was calculated for the comparison of renal damage between Stx2 and Control rats (C). Data are represented as mean ± SEM *P<0.05.

### Histological evaluation of renal damages

Kidneys obtained from Stx2 injected rats and Controls were microscopically evaluated. Histological evaluation of renal tissues from rats exposed to Stx2 showed glomerular mesangiolysis, tubular dilatation with loss of the brush border and widening of the tubular lumen as previously reported in mice HUS models [23]. Additionally, the percentage of tubular necrosis was evaluated. Stx2 treated rats showed a significant tubular necrosis compared with Control (**Figure 4**).

**Figure 4.**
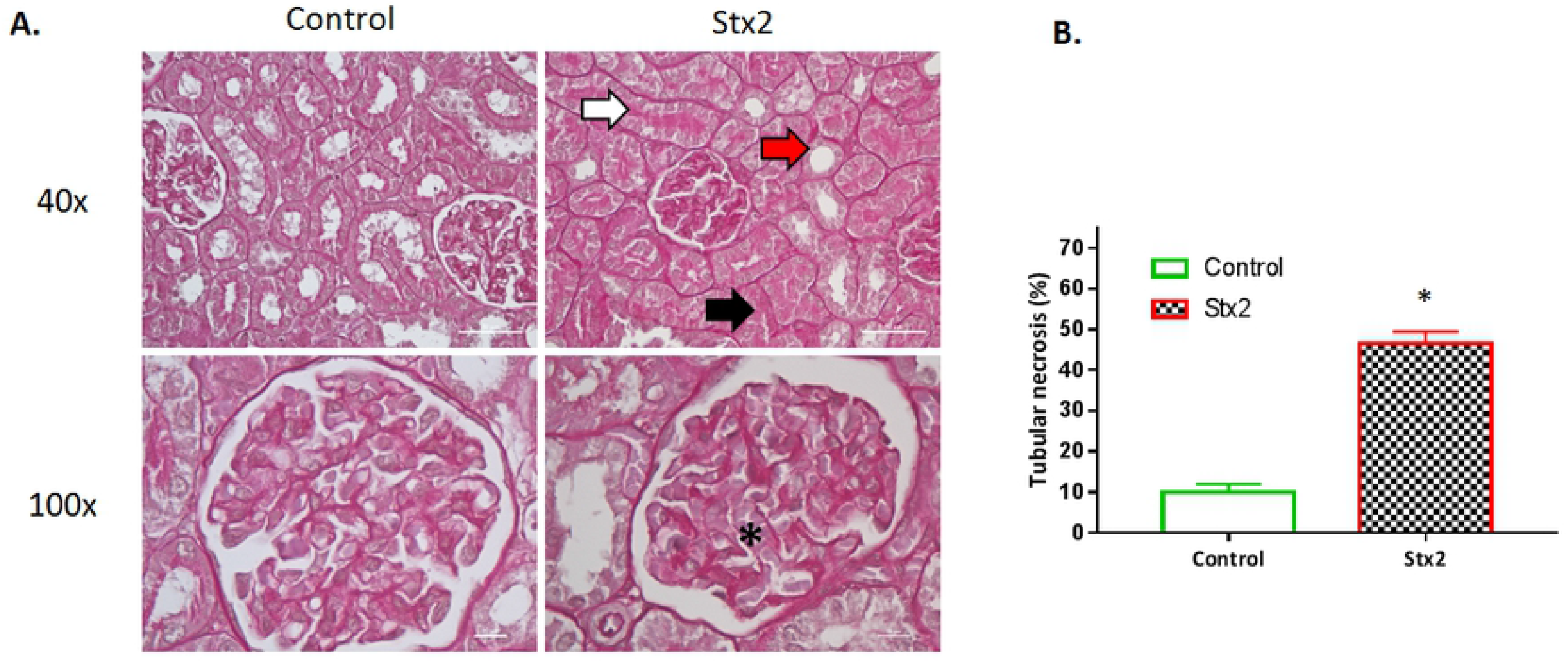
Histological evaluation of Stx2 damages in kidney tissues. Kidneys from Stx2 (n=6) and Control rats (n=4) were obtained after 48 h of injection. Then, were fixed in formol 10%, embedded in paraffin and sections of 5 μm were made using a microtome and mounted on 2% silane coated slides. The sections were stained with Periodic Acid-Schiff (PAS) and then observed and photographed under light microscopy. A representative image of Stx2 and PBS injected rats are shown. Glomerular mesangiolysis (asterisk) widening of the tubular lumen (red arrow) and intraluminal eosinophilic material (proteinaseus liquid) in tubes (withe arrow) are represented (A). Evaluation of percentage of tubular necrosis comparing Stx2 treated rats and Controls (B). Data are represented as mean ± SEM *P<0.05.

### Evaluation of the hydrodynamic diameter and size distribution of obtained MVs

DLS analysis of isolated MVs showed a bimodal size distribution. Briefly, a main fraction of 218.4 ± 56.2 nm (87.5%) and a smaller one of 17.2 ± 1.3 nm (17.1%) (mean ± SD) were observed. Further, the PDI value of 0.369 ± 0.026 nm (mean ± SD) was consistent with a bimodal size distribution.

### Total MVs and MVs-Stx2+ after toxin injection

Total MVs and MVs-Stx2+ from plasma of Stx2 treated and Control rats were analyzed by flow cytometry. No significant differences were obtained for total obtained MVs, at 48, 72 and 96 h post injection between Stx2 treated and non-treated rats (**Figure 5)**. On the other hand, a significant detection of circulating MVs-Stx2+ in Stx2 injected rats was observed at 96 h after toxin exposure (**Figure 6A**). **Figure 6B** shows a representative dot plot analysis of one experiment. In addition, Stx2 was detected when MVs were permeabilized with saponin. Stx2 was not detected in the surface of the MVs (without saponin permeabilization).

**Figure 5.**
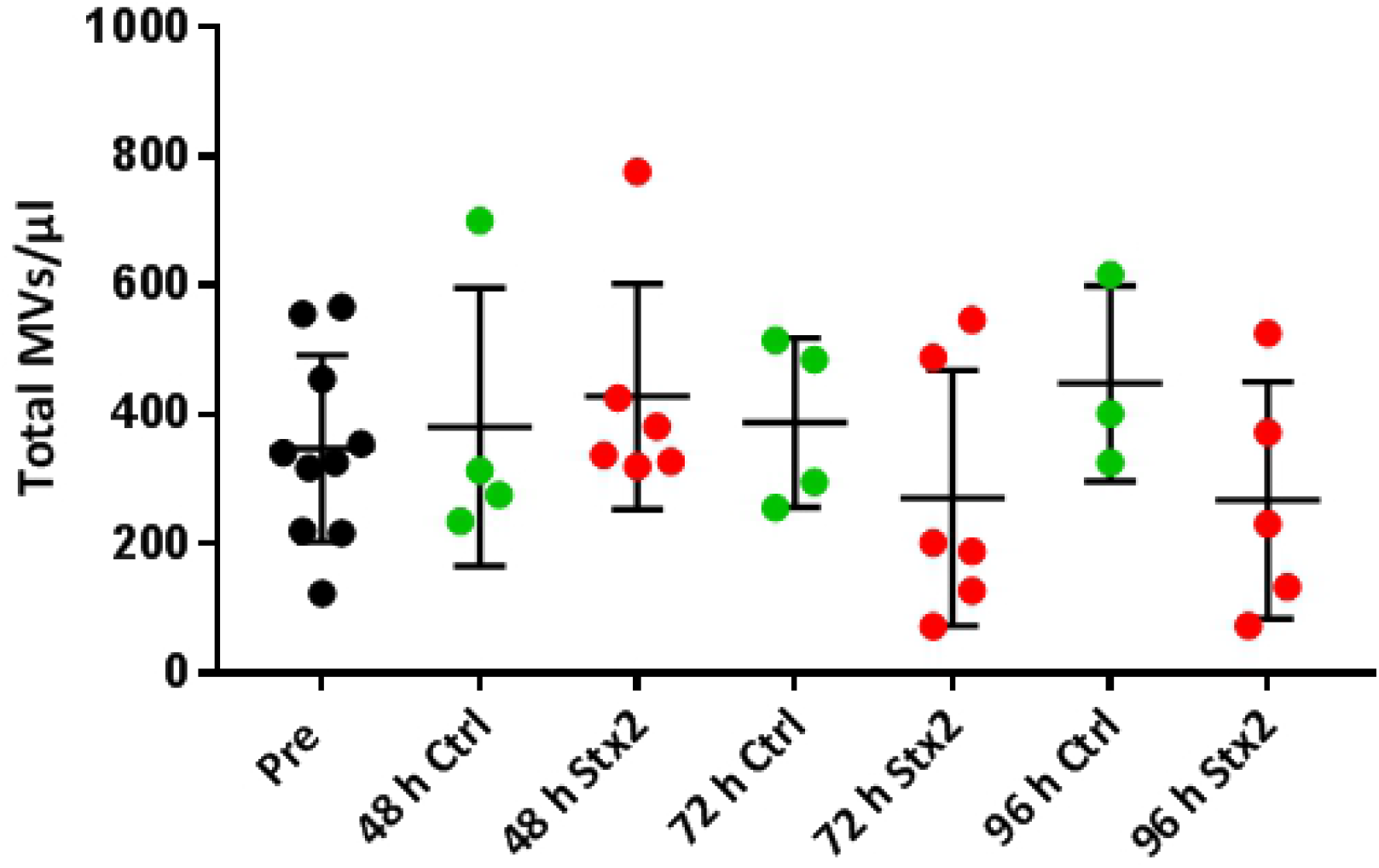
Evaluation of total circulating MVs after Stx2 injection in rats. Total MVs from plasma of Stx2 (n=5-6) and Control rats (n=3-4) at different time points after injection (48, 72 and 96 h) were obtained and labeled with AV-FITC dye and quantified (number of MVs/μl) by flow cytometry. Data are represented as mean ± SEM. P>0.05 n.s.

**Figure 6.**
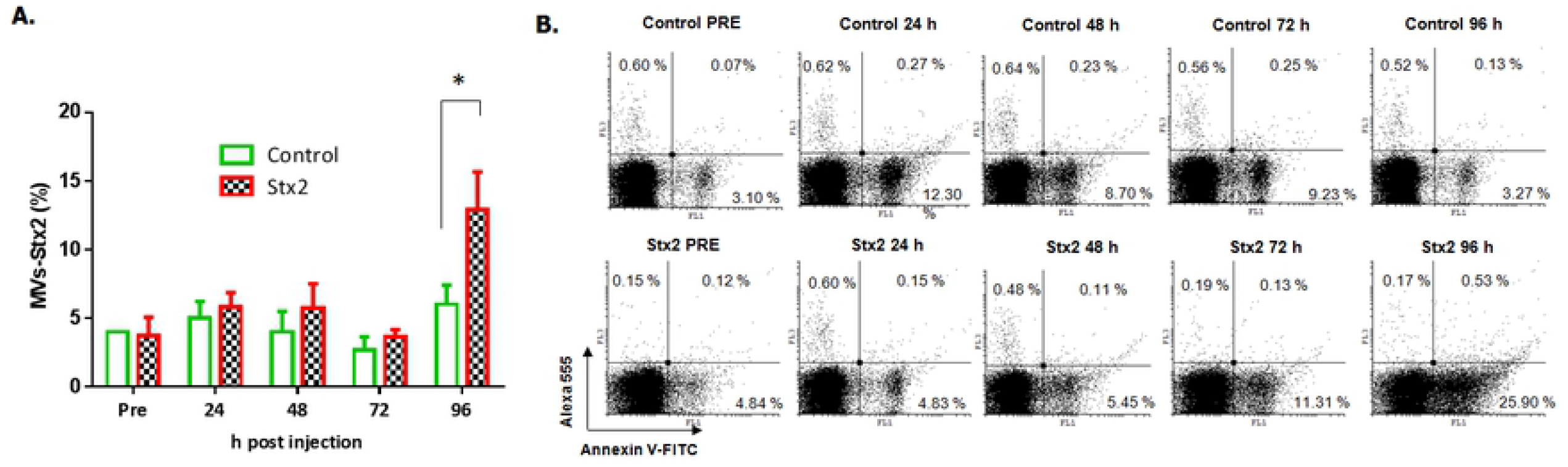
Stx2 positive MVs (MVs-Stx2+) after Stx2 injection. MVs-Stx2+ microvesicles in Stx2 (n=6) or Control rats (n=4) were detected by flow cytometry at different time points different time points after injection (Pre, 24, 48, 72, 96 h). The percentage of MVs-Stx2+ / total circulating MVs was calculated and compared between Stx2 treated rats and Control (A) Representative dot plots of each condition are shown (B) The experiment was repeated three times. Data are represented as mean ± SEM.*P<0.05.

### Correlation between plasma creatinine or urea and MVs-Stx2+

When plasma creatinine and urea values were correlated with the percentage of Stx2 positive microvesicles, no correlation was observed with both plasma parameters and circulating MVs containing Stx2 at 96 h after toxin injection. (**Figure 7**)

**Figure 7.**
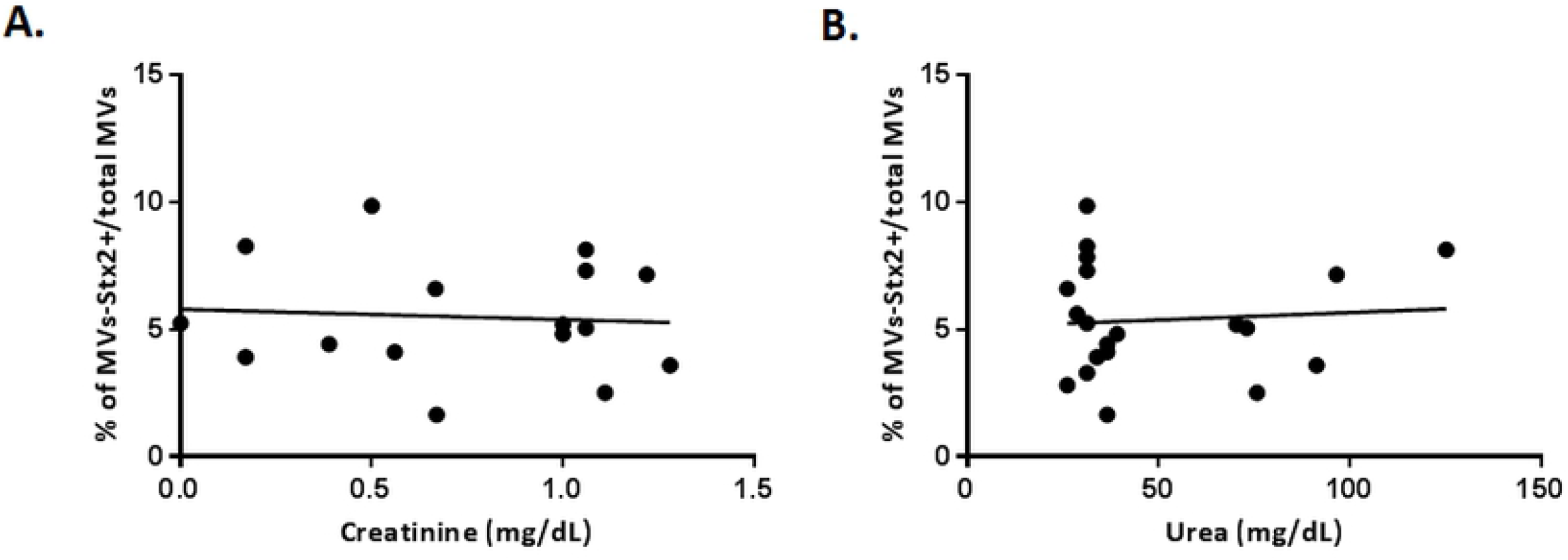
Correlation analysis between plasma creatinine or urea *vs* MV’s/Stx2+. Correlation analysis of MVs-Stx2+and creatinine (A) or urea (B) was evaluated at 96 h after Stx2 injection. Stx2: n=6 and PBS: n=4, P>0.05 n.s.

## DISCUSSION

Stx is the main virulence factor of STEC and is responsible for triggering HUS, clinically described by hemolytic anemia, thrombocytopenia and acute renal failure. Children under 5 years of age are the most affected by HUS, and Stx2 is the subtype toxin more frequently associated with severest cases of HUS. The early diagnosis of STEC infection is critical for supportive care and clinical management of the patient. Detection of Stx2 or STEC in fecal samples is used for routine testing, being in several cases time consuming, delaying the result. Detection of free toxin is minimal in blood circulation because transport of toxin to the target organs may be through binding blood cells or via MVs [24-26]. It was reported that MVs can be proinflammatory mediators and are elevated in HUS patients. In this regard, it has been also demonstrated that MVs can be responsible for coagulation disorders [27], and complement activation [28]. Altogether this data suggests that MVs may be also responsible for microangiopathic thrombosis characteristic of HUS.

It has been extensively described that Stx2 enters the endothelial cells by receptor mediated endocytosis. The globotriaosylceramide (Gb3) has been considered the principal receptor for Stx2. However, there is still some controversy about how Stx2 may affect non expressing Gb3 cells as intestinal cells [29] or neutrophiles [30]. In this sense, MVs-Stx2+ transfer may be an additional bacterial strategy to reach target cells with its virulence factors by a Gb3 dependent or independent manner [26]. Additionally, this strategy includes the evasion of the immune system, perpetuating bacterial survival. It has been reported that inflammatory and infection conditions may increase circulating MVs levels and that blood cell derived-MVs may play an important role in the development of HUS. Related to this, MVs-Stx2+ were proposed as a novel mechanism of toxin transfer between cells and as an alternative mechanism of toxin transport and cell damage [20]. According with these data, we propose that MVs-Stx2+ can be considered as a new biomarker for HUS diagnosis or STEC infection contributing to achieve the definitive diagnosis.

In this work, we analyzed the total circulating MVs and the MVs-Stx2+ in plasma samples from Stx2 injected rats and compared them with Controls (Stx2 non-treated animals, PBS injected). Firstly, we corroborated the detrimental effects of Stx2 as decrease of food ingestion and the consequently fall in the body weight. In addition, histological and functional renal alterations were also detected. These results were in accordance to our previous observations [31].

Of note, Stx2 injected rats exhibited the widely reported characteristics of body weight loss and food intake decrease, probably due to the general discomfort induced by cell damages of toxin in different target organs [23, 32]. Regarding the toxin effects on kidney, that is the main organ affected in HUS patients, we found a significant increase in serum creatinine and urea levels, thus indicating a renal dysfunction, corroborated with the decrease in the creatinine clearance. Thus, confirming the classical renal damages observed in HUS experimental animal models [33]. After that, we investigated total circulating MVs and MVs carrying Stx2 (MVs-Stx2+) by flow cytometry. MVs are spheroidal vesicles released from cells and they play an important role in cell–to–cell communication [34]. MVs are originated by an outward budding and fission of the plasma membrane and subsequently released into the extracellular space by ectocytosis. MVs range in size from 100 nm to 1,000 nm in diameter but can be even larger (up to 10 μm) in the case of oncosomes [35]. It has been widely accepted that MVs expose on their surface phosphatidilserine (PS), however, some evidence supports the notion that not all platelets derived MVs expose PS on their surface [36]. In our work, the size distribution of the isolated MVs showed a two size populations of approximately 160-280 nm and 16-18 nm. The most numerous particle population had the size of around 200 nm (almost 87 %). Probably the minority population sizing less than 20 nm represents cellular debris. However, this population was excluded in the flow cytometry analysis.

In our model, total MVs were quantified without characterization of their cell origin and the appearance of MVs-Stx2+ in plasma at different times after toxin injection was investigated. We found that the number of total MVs was not different between Stx2 injected rats and Control animals. In this regard, Stahl *et al*. quantified specific platelet or neutrophil derived MVs and found an increase of them in Stx2 treated mice. So, we speculate that in our model an increase of these specific populations of MVs may be also possible.

Additionally, we found that Stx2 treated rats exhibited significant circulating levels of MVs-Stx2+ after 96 h of toxin exposure. We also confirmed that Stx2 was localized inside MVs and not on the membrane surface. Accordingly with our results, Stahl *et al* [20] described the presence of MVs-Stx2 in mice after Stx2 injection and demonstrated Stx2 injected mice had lower circulating MVs-Stx2+ comparing with STEC infected mice. These results probably indicate that bacterial factors may contribute to proinflammatory stimulus or a continuous release of toxin that may be reflected in higher levels of circulating MVs-Stx2+ compare to the injection of one dose of purified Stx2.

Finally, we observed no correlation of plasma creatinine and urea levels that are considered parameters for the renal damages with the percentage MVs-Stx2+, indicating that these independent values may be complementary used for clinical evaluation.

In summary, MVs-Stx2+ may be considered as a new biomarker for HUS diagnosis that may help physicians for clinical supervision and for taking decisions on the management, treatment and follow up of the patient. However, clinical data of HUS patients and circulating MV-Stx2+ are needed to evaluate this hypothesis.

## Data availability

The data that support the findings of this study are available from the corresponding author upon reasonable request.

## Credit author contribution

The author contributions are as follows: conceived and designed the experiments: FS, FG, MM, DC, MMA performed the experiments: FS, FG, MM, DC, MMA; analyzed the data: FS, FG, CJ, MM, DC, MMA; contributed reagents/materials/analysis tools: FS, FG, CJ, MM, DC, CI, MMA; wrote the paper: FS, CJ, MM, DC, MMA. The authors declare no conflict of interest.

## Acknowledgements

We thank Tomas Lombardo from IDEHU, School of Pharmacy and Biochemistry, University of Buenos Aires, for guidance in the detection of particles by flow cytometry. We also thank Agostina Presta from IFIBIO-Houssay, UBA-CONICET, for developing histology slides.

